# Rapid sensing of L-leucine by human and murine hypothalamic neurons: neurochemical and mechanistic insights

**DOI:** 10.1101/244111

**Authors:** Nicholas Heeley, Peter Kirwan, Tamana Darwish, Marion Arnaud, Mark Evans, Florian Merkle, Frank Reimann, Fiona Gribble, Clemence Blouet

## Abstract

**Objective:** Dietary proteins are sensed by hypothalamic neurons and strongly influence multiple aspects of metabolic health, including appetite, weight gain and adiposity. However, little is known about the mechanisms by which hypothalamic neural circuits controlling behavior and metabolism sense protein availability. The aim of this study is to characterize how neurons from the mediobasal hypothalamus respond to a signal of protein availability: the amino acid L-leucine.

**Methods:** We used primary cultures of post-weaning murine mediobasal hypothalamic neurons, hypothalamic neurons derived from human induced pluripotent stem cells, and calcium imaging to characterize rapid neuronal responses to physiological changes in extracellular L-Leucine concentration.

**Results:** A neurochemically diverse subset of both mouse and human hypothalamic neurons responded rapidly to L-leucine. Consistent with L-leucine’s anorexigenic role, we found that 25% of mouse MBH POMC neurons were activated by L-leucine. 10% MBH NPY neurons were inhibited by L-leucine, and leucine rapidly reduced AGRP secretion, providing a mechanism for the rapid leucine-induced inhibition of foraging behaviour in rodents. Surprisingly, none of the candidate mechanisms previously implicated in hypothalamic leucine sensing (K_ATP_ channels, mTORC1 signaling, amino-acid decarboxylation) were involved in the acute activity changes produced by L-leucine. Instead, our data indicate that leucine-induced neuronal activation involves a plasma membrane Ca^2+^ channel, whereas leucine-induced neuronal inhibition is mediated by inhibition of a store-operated Ca^2+^ current.

**Conclusions:** A subset of neurons in the mediobasal hypothalamus rapidly respond to physiological changes in extracellular leucine concentration. Leucine can produce both increases and decreases in neuronal Ca^2+^ concentrations in a neurochemically-diverse group of neurons, including some POMC and NPY/AGRP neurons. Our data reveal that leucine can signal through novel mechanisms to rapidly affect neuronal activity.

**Highlights:**

- A neurochemically diverse group of mouse and human hypothalamic neurons rapidly sense and respond to L-leucine
- Leucine can produce neuronal activation or neuronal inhibition via distinct and novel Ca^2+^ signaling mechanisms.
- Leucine activates 25% ARH POMC neurons
- Leucine inhibits 10% ARH NPY/AGRP neurons and reduces AGRP secretion from fasted mediobasal hypothalamic slices

## 1. Introduction

Dietary proteins strongly influence metabolic health via their effect on appetite, weight gain and adiposity (1–3). Compelling evidence indicates that protein is the macronutrient that has the largest impact on energy intake, and that protein intake is tightly regulated independently of fat, carbohydrate and energy intake (1, 4). Several peripheral signals, including gut peptides and FGF21 (5–7), have been implicated in protein-induced satiety. However, none of these peptides specifically signal protein availability, suggesting that additional protein-specific mechanisms are involved to allow a tight, fat- and carbohydrate-independent regulation of protein intake.

Levels of the branched-chain amino acid L-leucine (Leucine) are likely to represent a physiological signal of protein availability in the control of appetite and metabolism. In humans and rodents, circulating leucine levels rapidly increase following the ingestion of a protein dense meal (8–10), and consumption of a protein-diluted diet chronically reduces serum leucine levels (11). It is well documented in the literature that leucine signals protein abundance to regulate key anabolic functions including muscle protein synthesis (12) and beta-cell insulin release (13) at least in part via mTORC1 signaling, a pathway for which leucine is the main activator (14). Importantly, protein sources with higher anabolic value (assessed by their ability to promote growth or activate TOR signaling) or rich in branched-chain amino acids produce increased satiety in flies (15), rodents (16) and humans (8), reinforcing the idea that blood leucine levels may convey information about the anabolic value of the meal to regulate appetite. In addition, leucine supplementation is sufficient to produce satiety in healthy humans (17).

Signals encoding protein availability, including leucine, likely act on the brain to control energy balance. The brain rapidly senses and responds to changes in circulating leucine levels during nutritional transitions (10). Appetite-regulating neurons in the arcuate nucleus of the hypothalamus (ARH) are well positioned to sense changes in leucine levels and engage downstream neuroendocrine and behavioral neural circuits. Located in the vicinity of the median eminence, they are directly exposed to circulating nutrient levels during energy deprivation (18), and project neural processes to the median eminence parenchyma, allowing constant monitoring of blood-borne signals (19). Local nanoinjection of leucine into the MBH produces satiety. Remarkably, this response includes a rapid delay in meal initiation and reduction of meal size, indicating that leucine can acutely alter the activity of orexigenic and anorexigenic circuits (10). However, the molecular, neurophysiological and neurochemical mechanisms implicated in this rapid change in feeding behavior remain poorly understood.

In this study, we aimed to characterize the neurophysiological responses of mediobasal hypothalamic neurons to physiological changes in extracellular leucine levels, test leucine responsiveness of POMC and non-POMC MBH neuronal subsets, and investigate the mechanisms involved in rapid leucine detection. In order to observe rapid, cell-autonomous responses to leucine in multiple species, we performed our studies in freshly dissociated mouse MBH neurons as well as human pluripotent stem cell (hPSC)-derived hypothalamic neurons.

## 2. Materials and methods

### 2.1. Mice

All mice were group-housed and maintained in individually ventilated cages with standard bedding and enrichment. Mice were housed in a temperature and humidity-controlled room on a 12-hour light/dark cycle with ad libitum access to water and standard laboratory chow diet unless otherwise stated. C57/Bl6J males were obtained from Charles River UK. NPY-GFP mice (Jackson Labs Stock No:006417) were genotyped with the following primers: 5’-TATGTGGACGGGGCAGAAGATCCAGG-3’; 5’-CCCAGCTCACAT ATTTATCTAGAG-3’; 5’-GGTGCGGTTGCCGTACTGGA-3’. POMC-GFP mice (Jackson Labs Stock No 009593) were genotyped using the following primers: 5’-AAGTTCATCTGCACCACC G-3’; 5’-TCCTTGAAGAAGATGGTG CG-3’. All studies were approved by the local Ethics Committee and conducted according to the UK Home Office Animals (Scientific Procedures) Act 1986.

### 2.2 Primary culture of post-weaning mediobasal hypothalamic neurons

Primary cultures of mediobasal hypothalamic neurons were prepared from 4 to 6-week old C57/Bl6J, POMC-EFGP and NPY-GFP males fasted overnight. Throughout the extraction and culture process, all media contained physiologically relevant concentrations of glucose and amino acids, based on concentrations measured in rat cerebrospinal fluid and microdialysis extracts (10, 20, 21). Mice were killed by cervical dislocation. Brains were extracted and rapidly placed into ice-cold extraction media (Suppl Table 1). Brain sections of 0.75 µm were cut using a McIlwain tissue chopper. The MBH was then dissected from coronal slices and placed in extraction media on ice. Tissue was transferred to papain (20 U/ml, Worthington, Lakewood, NJ, USA) pre-heated at 37°C and digested for 30 minutes at 37°C under agitation (Thermomixer, 500 rpm). After digestion, tissue extracts from 6 to 7 animals were pooled, transferred to a tube containing extraction media with 3.5 U/ml DNase I from bovine pancreas (Sigma) using a glass Pasteur pipette with a fire polished 1.5 mm opening, and triturated with pipettes with decreasing diameters (prepared according to Nagy et al., 2006). The trituration supernatant was gently loaded on top of a BSA gradient (4% BSA prepared in extraction media (pH 7.4) loaded on top of 8% BSA (pH 7.4)), spun for 5 minutes at 300 rcf, and the pellet was resuspended in culture media. 100 µl of resuspended cells were plated on the glass portion of 35 mm dishes (MatTek Corporation), coated with poly-lysine (0.1 mg/ml) using a 0.5 mm trituration pipette, inside a cloning cylinder (8 mm^2^, Sigma). Plates were placed in an incubator (37°C, 5% CO_2_) for 1 hour. After one hour, an additional 2ml culture media was added and the cloning cylinder removed. 4 to 6 culture dishes were prepared on each experimental day. Each culture dish was imaged once and represented our experimental unit.

For longer cell maintenance, 0.3 nM FGF2 (Sigma) was added to the culture media and media was changed every 2 days.

### 2.2 **Calcium Imaging** (adapted from (22))

Cells were loaded with 5 µM Fura 2 AM dye (Life Technologies) for 30 minutes, washed with aCSF (supplemented with amino acids and glucose, see Supplementary methods) and imaged using an inverted fluorescence microscope (Olympus IX71, Olympus, Southend on Sea, UK) with a 40X oil-immersion objective lens. GFP (to identify POMC or NPY cells) was excited at 488 nm and fura-2 at 340 nm and 380 nm using a monochromator (Cairn Research, Faversham, UK) and a 75 W xenon arc lamp, and emissions were recorded using an Orca ER camera (Hamamatsu, Welwyn Garden City, UK), a dichroic mirror and a 510 nm long pass filter. All images were collected on MetaFluor software (Molecular Devices, Wokingham, UK). The ratio of fura-2 emissions at 340 and 380 nm (340/380 ratio) was used to monitor changes in the intracellular calcium concentration. Solutions were perfused continuously at a rate of approximately 0.5 ml/ minutes.

### 2.3 Immunofluorescent staining

Cells were fixed with 4% PFA, washed 3 times in 0.1% PBST, followed by blocking in 5% normal donkey serum in 0.3% PBST. Primary antibody, mouse anti – NeuN (1:250, Millipore), chicken anti-MAP2 (1:500, Abcam) or rabbit anti-Ser235/236 rpS6 (1:1000, Cell Signaling Technologies) was applied overnight at 4°C. After PBS wash, secondary antibody in 5% normal donkey serum, was applied for 1 hour. After PBS wash, Vectashield with DAPI (Vector Labs) was added to coverslips inverted onto SuperFrost Plus slides. We analyzed fluorescence colocalisation to evaluate the number of neurons using confocal microscopy (Zeiss LSM 510 Meta).

### 2.4 Trypan blue staining

Trypan Blue staining was performed 24 hours post plating. Trypan blue (0.2%, filtered, Sigma) was added to each well for 15 minutes followed, was washed 3 times with PBS. Cells were visualized by Brightfield microscopy on an Evos XL Core Cell Imaging System (Life Technologies). Images were processed using ImageJ and counted to assess the number of live neurons, dead neurons, and the amount of debris.

### 2.5 Human derived hypothalamic neurons

Human pluripotent stem cells (hPSCs) (13B iPSC) were maintained in mTESR1 media [Ludwig, 2006 #1095] and differentiated to hypothalamic neurons as previously described (23, 24). Human hypothalamic neurons were used at 30–50 days in vitro, at which point there were abundant neurons immunoreactive for POMC. To study cell-autonomous responsiveness to L-leucine, cultures were enzymatically dissociated with TrypLE and Papain as described previously (Kirwan, Jura et al. 2017) and replated onto glass-bottomed 35 mm dishes (MatTek Corporation) coated with Geltrex (Thermo Fisher Scientific, 21041025). Cells were loaded with 5 µM Fura 2 AM for 45 minutes and treated with vehicle or experimental solutions as described above.

### 2.6 AGRP secretion from medial hypothalamic slices

The release of AGRP from mouse hypothalamic slices was measured ex vivo as described by Enriori *et al*. (25). Extraction and culture media described in Supplementary methods and supplemented with 0.6 TIU aprotinin/ml were used. 2 mm medial hypothalamic slices containing the PVH and ARH were prepared from C57/Bl6J mice in ice-cold extraction media. Slices were equilibrated for 1 hour at 37C in culture media. Culture supernatant was collected following a 45 minutes with incubation in 300 µl culture media, a 45 minutes incubation with culture media + 500 µM Leu, and a final 45 minutes incubation with culture media + 56 mM KCL. AGRP levels were measured using an ELISA kit from Phoenix Pharmaceuticals.

### 2.7 Drugs Used

Thapsigargin (Sigma), Rapamycin (Millipore), Ryanodine (Insight Biotechnology) and 2-APB (Abcam) were prepared as 10000X stocks in DMSO. Ryanodine (Insight Biotechnology) was prepared as a 1000X stock in DMSO. Tolbumatide and Diazoxide (Sigma) were prepared as 200X stocks in 1 M Sodium Hydroxide, which was neutralized in final solution using an equivalent volume of 1 M HCl. BCH (2-amino-2-nobornanecarboxylic acid, Sigma) was prepared in the aCSF solution used for imaging. L-leucine, KIC and L-valine (Sigma) were prepared as 100X stocks in water.

### 2.8 Data and Statistical Analysis

Treatments were randomly assigned to each culture dish and relevant control conditions were systematically included on each experimental day. For each culture dish (experimental unit), imaging data from all cells within the 40X objective visual field were analyzed using standardized criteria for all our imaging experiments, allowing unbiased analysis. Data were smoothed using the sliding window method over 20 seconds. From these smoothed data, we calculated for each control and treatment periods 1- the area under the curve (AUC) over 8 minutes for 10 minutes exposure periods, excluding the first and last minute of each period, or over 4 minutes for 5 minutes exposure periods, excluding the first and last 30 seconds of each period, 2- the maximum Fura2 340/380 ratio, 3- the minimum Fura2 340/380 ratio and 4- the mean Fura2 340/380 ratio over 4 minutes or 8 minutes. Only cells showing a calcium response to KCl (max Fura2 340/380 ratio during KCl treatment at least 50% higher than baseline Fura2 340/380 ratio) and with a Fura2 340/380 ratio stable throughout baseline and washout periods (average Fura2 340/380 ratio during washout maximum 10% higher or lower than average Fura2 340/380 ratio at baseline) were included in the analysis. Cells were considered activated if showing a reversible increase in Fura2 340/380 ratio during the treatment period, with AUC of the Fura2 340/380 ratio during the treatment period at least 15% higher than during both vehicle treatment periods. Cells were considered inhibited if showing a reversible decrease in Fura2 340/380 ratio during the treatment period, with the AUC of the Fura2 340/380 ratio during the treatment period at least 10% lower than during both vehicle treatment periods. Traces were visually inspected to confirm the validity of the analysis.

All data, presented as means ± SEM, have been analyzed using GraphPad Prism 6. For all statistical tests, an α risk of 5% was used. All dataset with multiple treatments were analyzed using repeated-measures two-way ANOVAs and adjusted with Bonferroni’s post hoc tests. Multiple comparisons were tested with one-way ANOVAs and adjusted with Tukey’s post hoc tests. Single comparisons were made using one-tail Student’s t tests.

## Supplementary Methods

### Preparation of extraction and culture media

Media were prepared as described in Supplementary Table 1 (without amino acids, glucose/pyruvate/lactate or B27), sterile filtered, stored at 4°C and used for up to 6 months. Osmolarity was adjusted to 260 mOsm using sucrose. The day before culturing cells, final extraction and culture media were prepared by adding B27 without Insulin (Life Technologies), Amino acid stock solutions (Supplementary Table 2) and glucose, pyruvate, and lactate (final concentrations: Glucose, 2.5 mM; Lactic Acid, 1 µM; Sodium Pyruvate, 0.23 mM). Papain (40 U/ml, PAP2, Worthington Biochemical) was prepared in extraction media (with amino acids, glucose, pyruvate and lactate, as above, but without calcium and B27) and diluted to a final concentration of 20 U/ml. Papain was not used for more than one week after reconstitution. aCSF was prepared as described in Supplementary Table 3), with amino acids added to the same concentration as in extraction and culture media (see Appendix 1), and 2.5 mM glucose.

**Supplementary Table S1:** Vitamin and Salt composition of culture and extraction media

| Components | Culture media (mM) | Extraction media (mM) |
| --- | --- | --- |
| **Vitamins** |  |  |
| Choline chloride | 0.029 | 0.029 |
| D-Calcium pantothenate | 0.008 | 0.008 |
| Folic Acid | 0.009 | 0.009 |
| Niacinamide | 0.033 | 0.033 |
| Pyridoxal hydrochloride | 0.020 | 0.020 |
| Riboflavin | 0.001 | 0.001 |
| Thiamine hydrochloride | 0.012 | 0.012 |
| Vita minutes B12 | 5.02E-06 | 5.02E-06 |
| i-Inositol | 0.040 | 0.040 |
| **Inorganic Salts** |  |  |
| Calcium Chloride | 1.801 | 1.801 |
| Ferric Nitrate | 2.48E-04 | 2.48E-04 |
| Magnesium Chloride | 0.814 | 0.814 |
| Potassium Chloride | 5.333 | 5.333 |
| Sodium Bicarbonate | 26.190 | 0.88 |
| Sodium Chloride | 51.724 | 76 |
| Sodium Phosphate monobasic | 0.906 | 0.906 |
| Zinc sulfate | 6.74E-04 | 6.74E-04 |
| **Other Components** |  |  |
| HEPES | 10.924 |  |
| Phenol Red | 0.022 | 0.0212 |
| MOPS |  | 10 |
| pH and Temperature | pH 7.6 at 37°C | pH 7.4 at 20°C |

**Supplementary Table S2:**
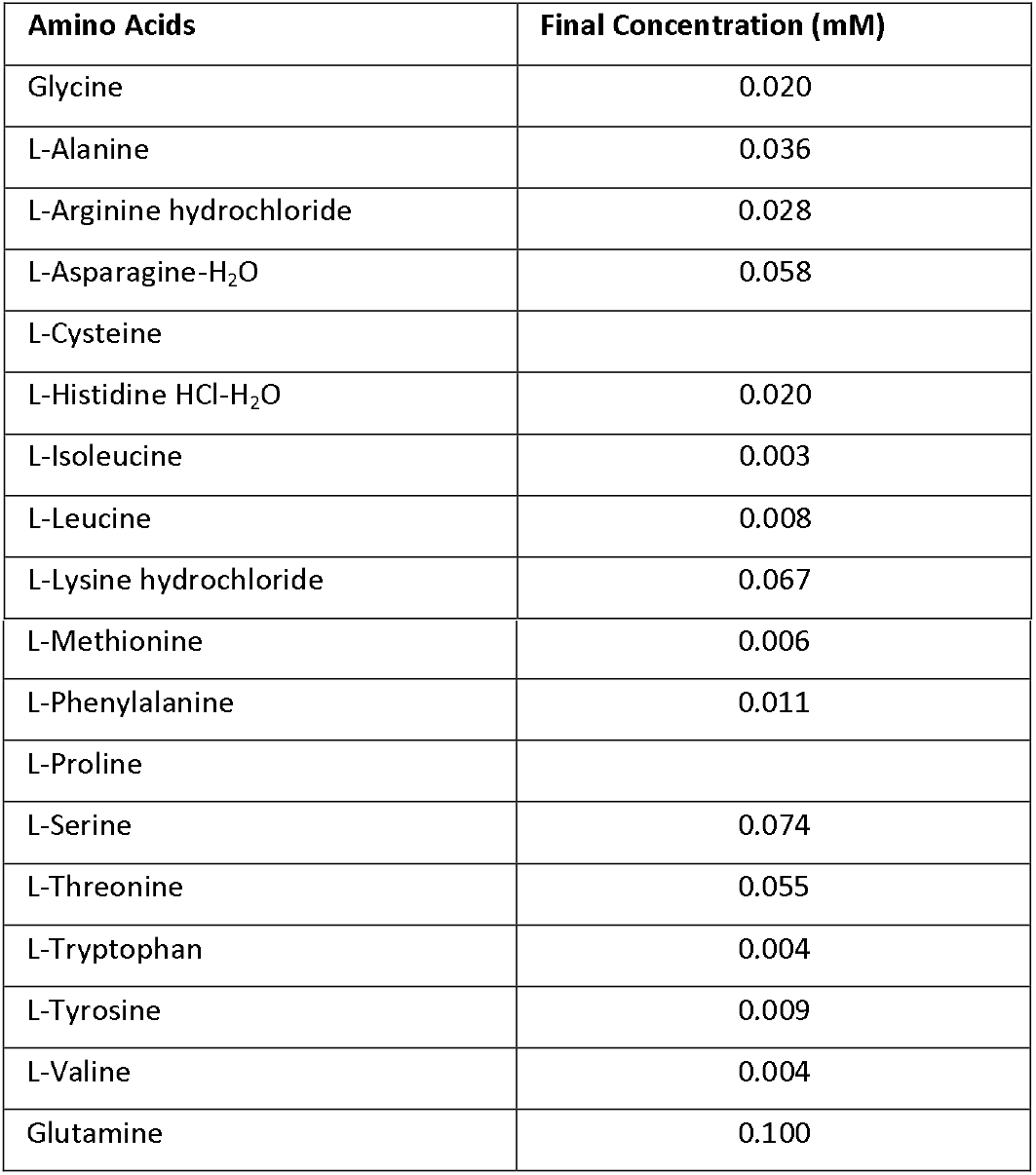
Amino acid composition of extraction and culture media

**Supplementary Table S3:** aCSF salt composition (pH adjusted to 7.4).

|  | Concentration (mM) |
| --- | --- |
| NaCl | 138 |
| KCl | 4.5 |
| NaHCO<sub>3</sub> | 4.2 |
| NaH<sub>2</sub>PO<sub>4</sub> | 1.2 |
| HEPES | 10 |
| CaCl<sub>2</sub> | 2.6 |
| MgCl<sub>2</sub> | 1.2 |

## 3 Results

### 3.1 Culture of post-weaning mouse mediobasal hypothalamic neurons

To characterize the acute cell-autonomous neurophysiological responses of murine mediobasal hypothalamic neurons to physiological changes in extracellular leucine levels and investigate the molecular and neurochemical underpinnings, we first optimized calcium imaging of freshly dissociated mouse mediobasal hypothalamic (MBH) neurons. We used mice between 4- to 6-weeks of age to investigate leucine-sensing properties of post-weaning MBH neurons. We optimized a culture protocol based on the published literature (26, 27) (Figure 1a) (Supplementary Figure 1). Commercially available neuronal culture media contain supra-physiological concentrations of glucose and amino acids, preventing the study of how physiological changes in nutrient concentrations are sensed by neurons. We therefore produced our own extraction and culture media with physiological nutrient concentrations based on those measured in rat cerebrospinal fluid and hypothalamic microdialysis extracts (see Material and Methods) (10, 20). These media did not affect cellular yield and viability (assessed by trypan blue staining on neurons up to 3 days in culture) compared to cultures prepared with Hibernate A (BrainBits) and Neurobasal A (Invitrogen) (data not shown).

**Figure 1:**
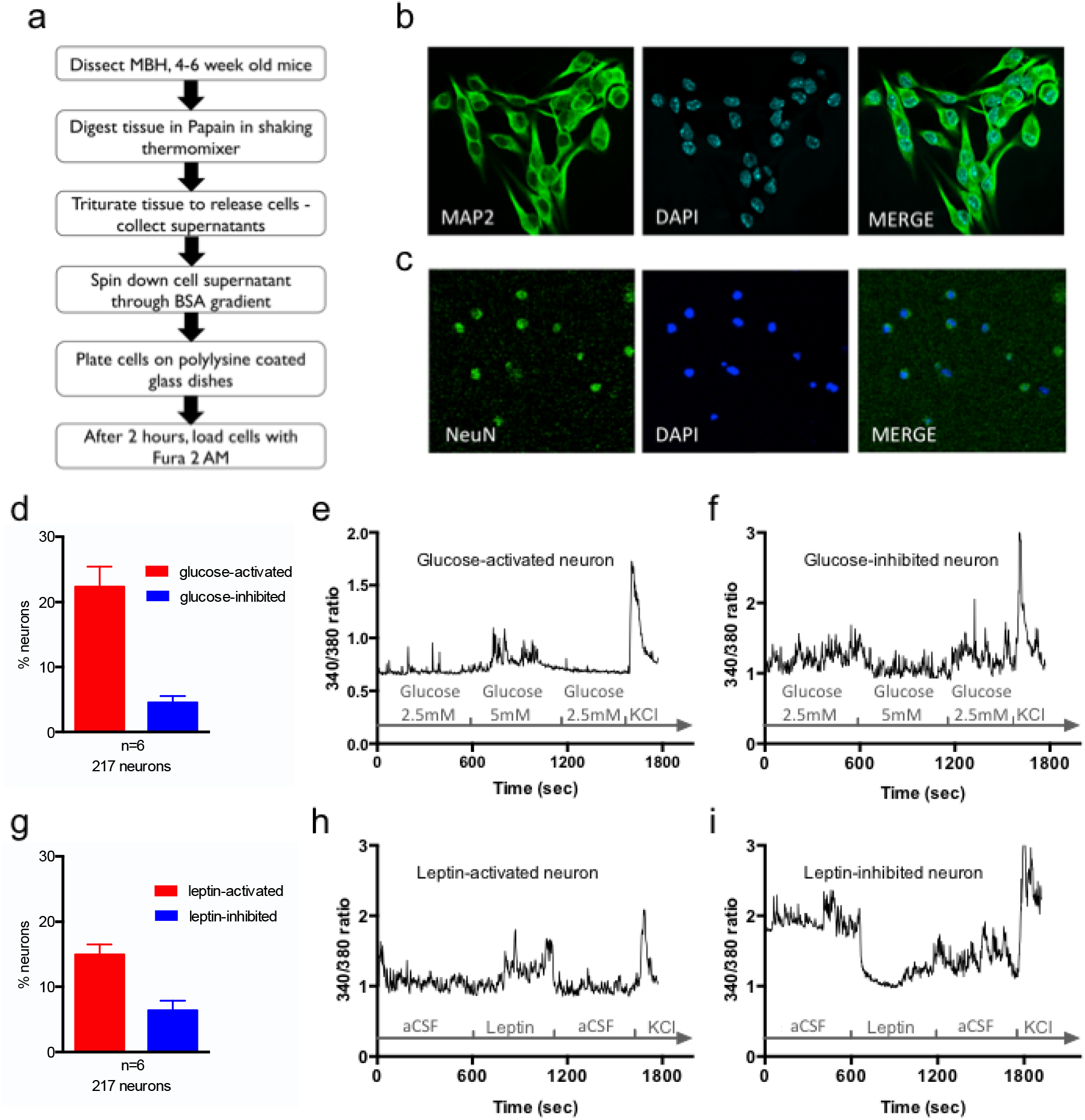
Primary culture of dissociated post-weaning mediobasal hypothalamic neurons sense glucose and leptin. Protocol to prepare dissociated mouse mediobasal hypothalamic neurons for primary culture and calcium imaging (a). MAP2 (b) and NeuN (c) immunostaining in primary culture of mediobasal hypothalamic neurons. Glucose-induced (d, e, f) and leptin-induced (g, h, i) calcium responses, assessed via 340/380nm fluorescence ratios (reflecting [Ca^2+^]_i_), in mouse mediobasal hypothalamic neurons in culture exposed to a transition from 2.5mM to 5mM glucose or to 10nM leptin. Data are means ± SEM.

As expected (27), cells in culture were predominantly neuronal and expressed MAP2 and NeuN (Figure 1b and 1c). We assessed the responsiveness of these cultures to glucose and leptin, two metabolic signals sensed by mediobasal hypothalamic neurons. We found that a transition from 2.5mM to 5mM glucose triggered an increase in intracellular Ca^2+^ concentration [Ca^2+^]_i_ in 25% of neurons in culture, and a decrease in [Ca^2+^]_i_ in 4.75% of neurons (Figure 1d-1f), consistent with previous observations (28, 29). 10nM Leptin increased [Ca^2+^]_i_ in 15% of neurons, and decreased [Ca^2+^]_i_ in 6.7% of neurons, consistent with previous reports (Figure 1g-1i) (30). These data indicate that under these culture conditions, dissociated MBH neurons from post-weaning mice can survive and maintain metabolic sensing properties.

### 3.2 Leucine rapidly alters [Ca^2+^]_i_ in a subset of mediobasal hypothalamic neurons

We then went on to assess changes in [Ca^2+^]_i_ in response to a 10 minutes exposure to 500µM leucine, a concentration consistent with the physiological range of postprandial circulating leucine levels (Blouet, 2009). Cells were exposed to a custom-made aCSF media containing low nutrient levels (see Supplementary methods) during the first 10 minutes baseline period as well as during the 10–20 minutes treatment period in control wells exposed to vehicle only. Treatment with aCSF induced calcium responses in a small fraction of cells (<3%, comparing [Ca^2+^]_i_ following treatment with aCSF compared to the baseline period)(Figure 2a). Treatment with leucine produced elevated [Ca^2+^]_i_ in 9% of neurons in culture (leucine-activated neurons) and a decrease in [Ca^2+^]_i_ in 14% of neurons in culture (leucine-inhibited neurons) (Figure 2a, 2b, 2c). Leucine-inhibited neurons were active during the baseline low-nutrient condition period, as indicated by the higher baseline [Ca^2+^]_i_ during vehicle exposure compared to leucine-activated neurons (Figure 2b and 2c, 2e and 2g). In activated cells, leucine produced a peak in the Fura2 340/380 ratio 46% above that seen in vehicle treated cells, and a 48% increase in the Fura2 340/380 AUC (Figure 2d-2e). In inhibited cells, leucine produced an average 61% decrease in Fura2 340/380 AUC (Figure 2f-2g). These data indicate that leucine can activate or inhibit subsets of mediobasal hypothalamic neurons and that a total of 23% of mediobasal hypothalamic neurons are primary leucine sensors. By contrast, 500µM valine activated only 2.7% and inhibited 0.4% of neurons in culture (n=4, 225 neurons).

**Figure 2:**
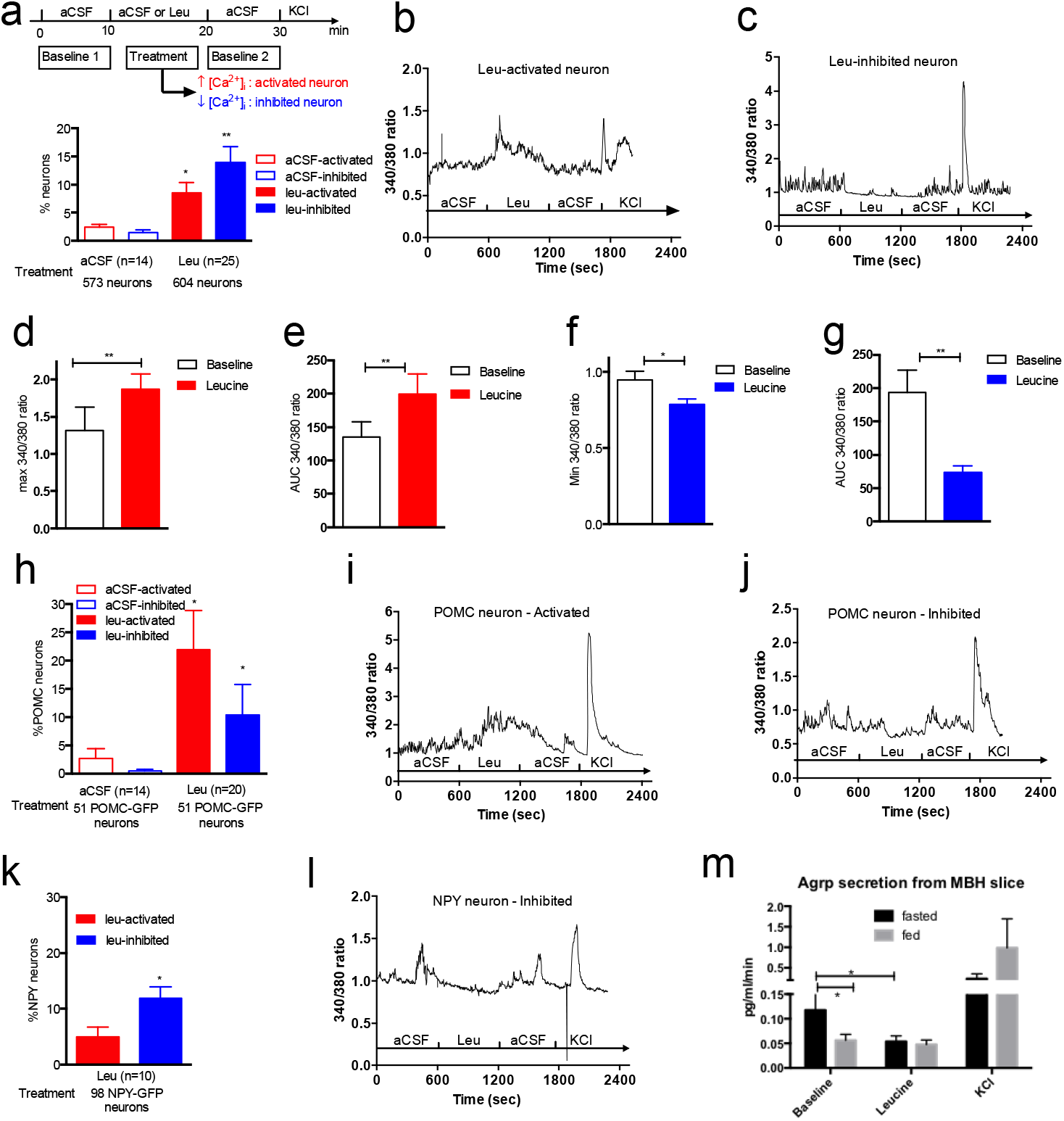
Leucine rapidly alters [Ca^2+^]_i_ in a neurochemically-diverse subset of mouse mediobasal hypothalamic neurons. Leucine-induced changes in [Ca^2+^]_i_ (a, b, c) as assessed by 340/380nm fluorescence ratios in primary cultures of mediobasal hypothalamic neurons. Peak 340/380nm ratio (d) and 340/380nm ratio AUC (e) during aCSF or leucine exposure in leucine-activated neurons. Minimum 340/380nm ratio (f) and 340/380nm ratio AUC (g) during aCSF or leucine exposure in leucine-inhibited neurons. Calcium responses to leucine in primary cultures of mediobasal hypothalamic POMC-GFP neurons (h). Leucine-induced changes in [Ca^2+^]_i_ (i, j) as assessed by 340/380nm fluorescence ratios in POMC-GFP neurons in culture. Calcium responses to leucine in primary cultures of mediobasal hypothalamic NPY-GFP neurons (k). Leucine-induced changes in [Ca^2+^]_I_ (l) as assessed by 340/380nm fluorescence ratios in NPY-GFP neurons in culture. AGRP secretion from 1mm medial hypothalamic slices from fasted (n=11) or fed (n=10) animals exposed to leucine. Data are means ± SEM. *:p<0.05. **: p<0.01

### 3.3 Leucine is rapidly sensed by MBH POMC and NPY neurons

We characterized leucine responses of MBH POMC and NPY neurons using cultures obtained from POMC-EGFP and NPY-hrGFP mice. Leucine activated 22% of POMC-EGFP neurons and inhibited 10% of POMC-EGFP neurons (Fig 2h-2j). By contrast, only a small fraction of NPY-GFP neurons were activated by leucine and 14% of NPY-GFP neurons were inhibited by leucine (Fig 2k-2l). To further establish the ability of leucine to inhibit ARH NPY/AGRP neurons, we measured AGRP secretion from hypothalamic sections maintained ex vivo with or without leucine. Leucine failed to alter AGRP secretion from sections obtained from fed animals. By contract, leucine significantly suppressed AGRP secretion from hypothalamic sections obtained from fasted animals (Figure 2m). Thus, subsets of POMC and NPY neurons are leucine responsive, and while leucine can either activate or inhibit these 2 neuronal subpopulations, a majority of leucine-sensing POMC neurons are activated by increased leucine concentrations, and a majority of leucine-sensing NPY neurons are inhibited by increased leucine concentrations.

### 3.4 Rapid sensing of Leucine by MBH neurons is independent of mTORC1 signaling.

The mTORC1 signaling pathway is the best-characterized intracellular leucine sensing pathway and has been implicated in MBH leucine sensing and the anorectic effects of hypothalamic leucine exposure [Cota, 2006 #17] [Blouet, 2008 #15], Treatment of mouse MBH primary cultures with 10nM rapamycin for 1h suppressed S6 ribosomal Ser240/244 phosphorylation, a surrogate marker for mTORC1 activity (Figure 3a). Under these conditions, we found that leucine was still able to produce neuronal activation and inhibition, at frequencies similar to those measured during exposure to vehicle control (Figure 3b-3d). Thus, rapid changes in [Ca^2+^]_i_ in response to leucine in MBH neurons do not rely on activation of mTORC1.

**Figure 3:**
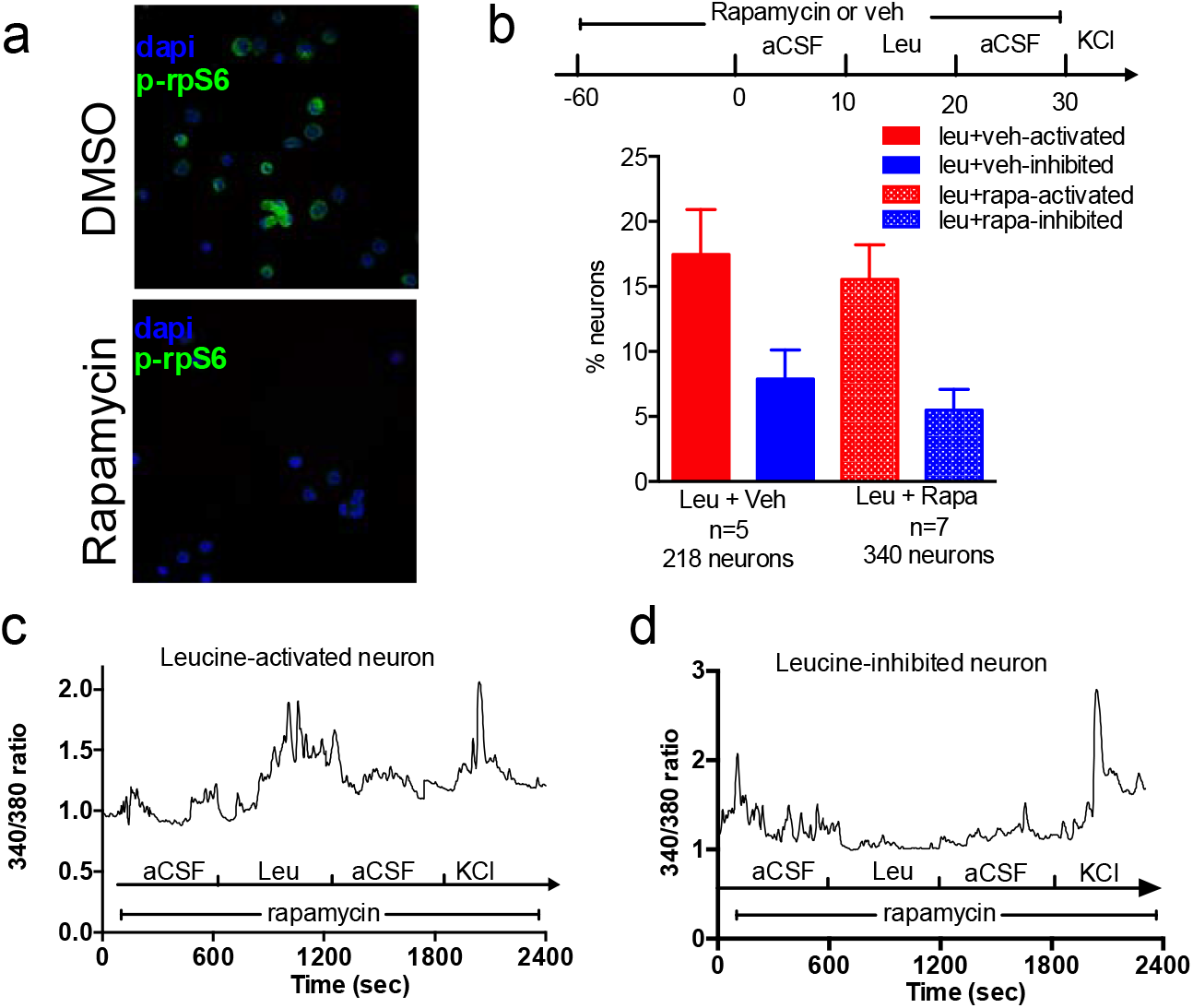
Leucine-induced changes in [Ca^2+^]_i_ are independent of mTORC1 signaling. Immunostaining against ribosomal protein S6 phosphorylated at Ser 240/244 (p-rpS6) in primary cultures of mediobasal hypothalamic neurons treated for 1h with 10nM rapamycin (a). Neuronal responses to leucine in primary cultures of mediobasal hypothalamic neurons pre-treated with rapamycin (b). Leucine-induced changes in [Ca^2+^]_i_ as assessed by 340/380nm fluorescence ratios in primary cultures of mediobasal hypothalamic neurons treated with rapamycin (c, d). Data are means ± SEM.

### 3.5 Rapid sensing of Leucine by MBH neurons is independent of leucine intracellular metabolism and K_ATP_ channels.

ATP production and closure of K_ATP_ channels have been implicated in nutrient sensing in pancreatic beta-cells and activation of mediobasal hypothalamic neurons sensitive to glucose and oleic-acid [Beall, #400; Jo, 2009 #401]. Therefore, we tested the role of this pathway in leucine-induced neuronal activation using a set of complementary approaches. We reasoned that leucine, via its catabolism to alpha-ketoisocaproic acid (KIC), promotes the production of TCA cycle intermediates, eventually leading to increases in intracellular ATP levels, decreases in intracellular MgADP levels, and potential closure of K_ATP_ channels which could contribute to leucine-induced neuronal activation.

We first tested the effect of KIC, leucine’s first catabolite, on neuronal activation in our preparation. Treatment with isomolar concentrations of KIC produced an increase in [Ca^2+^]_i_ in only 4% of neurons, and a decrease in [Ca^2+^]_i_ in only 3% of neurons (Figure 4a), indicating that leucine’s metabolite KIC cannot replicate leucine’s rapid effects on hypothalamic neuronal activity.

**Figure 4.**
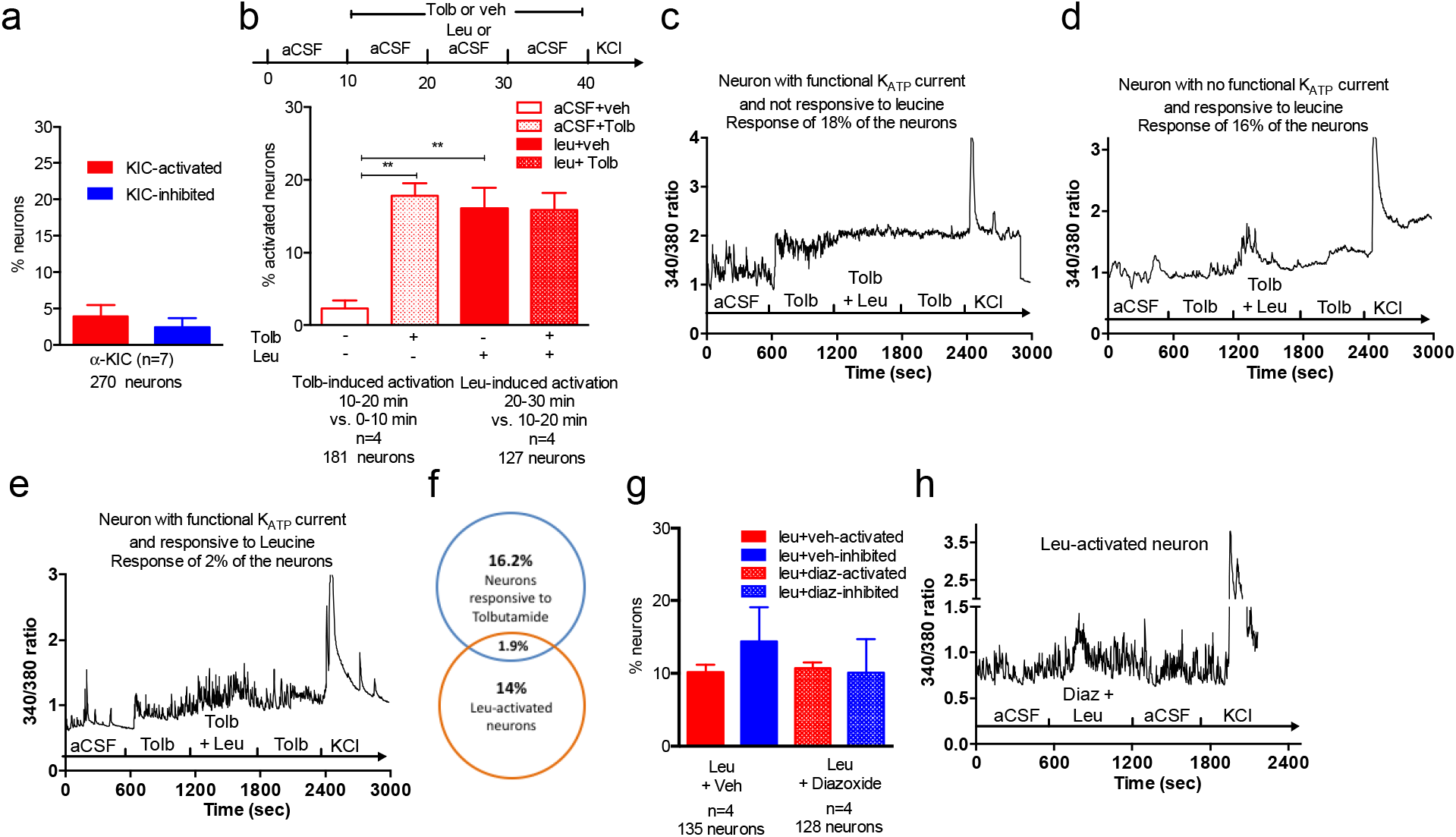
Leucine-induced changes in [Ca^2+^]_i_ are independent of leucine metabolism and K_ATP_ channels. Changes in [Ca^2+^]_i_ in primary cultures of mediobasal hypothalamic neurons treated with KIC (a), leucine in the presence of Tolbutamide (b, c, d, e, f), leucine in the presence of diazoxide (diaz) (g,h) Data are means ± SEM. *:p<0.05. **: p<0.01

We then used the K_ATP_ channel blocker tolbutamide and the K_ATP_ channel opener diazoxide to directly interrogate the role of K_ATP_ Channels in MBH leucine sensing. Tolbutamide blocks hypothalamic K_ATP_ channels at doses of 100 µM and above (31, 32). Pre-treatment with 500 µM Tolbutamide produced an increase in [Ca^2+^]_i_ compared to [Ca^2+^]_i_ during the baseline aCSF exposure period in 18% of the total neuronal population in culture (Figure 4b-4c) and 19% of POMC neurons. This result is consistent with previous findings indicating that 20% of MBH neurons express K_ATP_ channels (28). During co-administration of leucine and tolbutamide in the next 10 minutes of the recording session, 16% of cells exhibited an increased [Ca^2+^]_i_ compared with levels recorded during the tolbutamide only exposure (Figure 4b and 4d). Only 2% of cells responded to both tolbutamide and leucine (Figure 4e). Under these conditions, 25% of POMC neurons were activated by leucine but none of these leucine-activated POMC neurons were activated by tolbutamide alone. These results indicate that a majority of leucine-activated cells have no functional K_ATP_ currents, and closure of K_ATP_ channels does not alter the ability of MBH neurons to rapidly sense leucine. We then tested the effect of the K_ATP_ channel opener diazoxide. At concentrations around 300/400 µM, diazoxide hyperpolarizes cells expressing K_ATP_ channels within 2 minutes (33, 34). Co-administration of 340 µM diazoxide with leucine failed to abrogate leucine-induced changes in [Ca^2+^]_i_ (Fig 4g-4h). Collectively these data indicate that leucine intracellular metabolism and closure of K_ATP_ channels do not account for the rapid sensing of leucine by MBH neurons.

### 3.6 Extracellular leucine sensing mediates rapid leucine detection by MBH neurons

Leucine is transported across the cell membrane principally via system L amino acid transporters LAT1 (slc7a5) and LAT2 (slc7a8) (35), both expressed in hypothalamic POMC and AGRP neurons (36). We tested the role of system L transporter in MBH leucine sensing using 2-aminobicyclo-(2,2,1)-heptane-2-carboxylic acid (BCH), a well-characterized inhibitor of system L amino acid transporters (37). BCH is a non-metabolisable analogue of Leucine that saturates all members of LAT family at 10mM. Of note, in pancreatic beta-cells, BCH stimulates insulin secretion via activation of glutamate dehydrogenase (GDH), anaplerosis, replenishment of TCA cycle intermediates, ATP production and closure of K_ATP_ channels [Han, 2016 #1252], Therefore, we first examined the effect of BCH on its own on [Ca^2+^]_i_ in our primary culture model.

Pre-treatment with 10mM BCH alone activated 10% of neurons in culture (Figure 5a and 5b). During co-administration of leucine and BCH during the next 10 minutes of the recording session, 12% of neurons showed an increase in [Ca^2+^]_i_ compared to [Ca^2+^]_i_ during the BCH only exposure (Figure 5c). Only 1% of cells showed a response to both BCH and leucine (Figure 5d). Likewise, inhibition of leucine transport failed to abrogate leucine-induced inhibition of MBH neurons (Figure 5e and 5f). These results indicate that leucine sensing is independent of system L transporters, and suggest that leucine may be sensed extracellularly.

**Figure 5.**
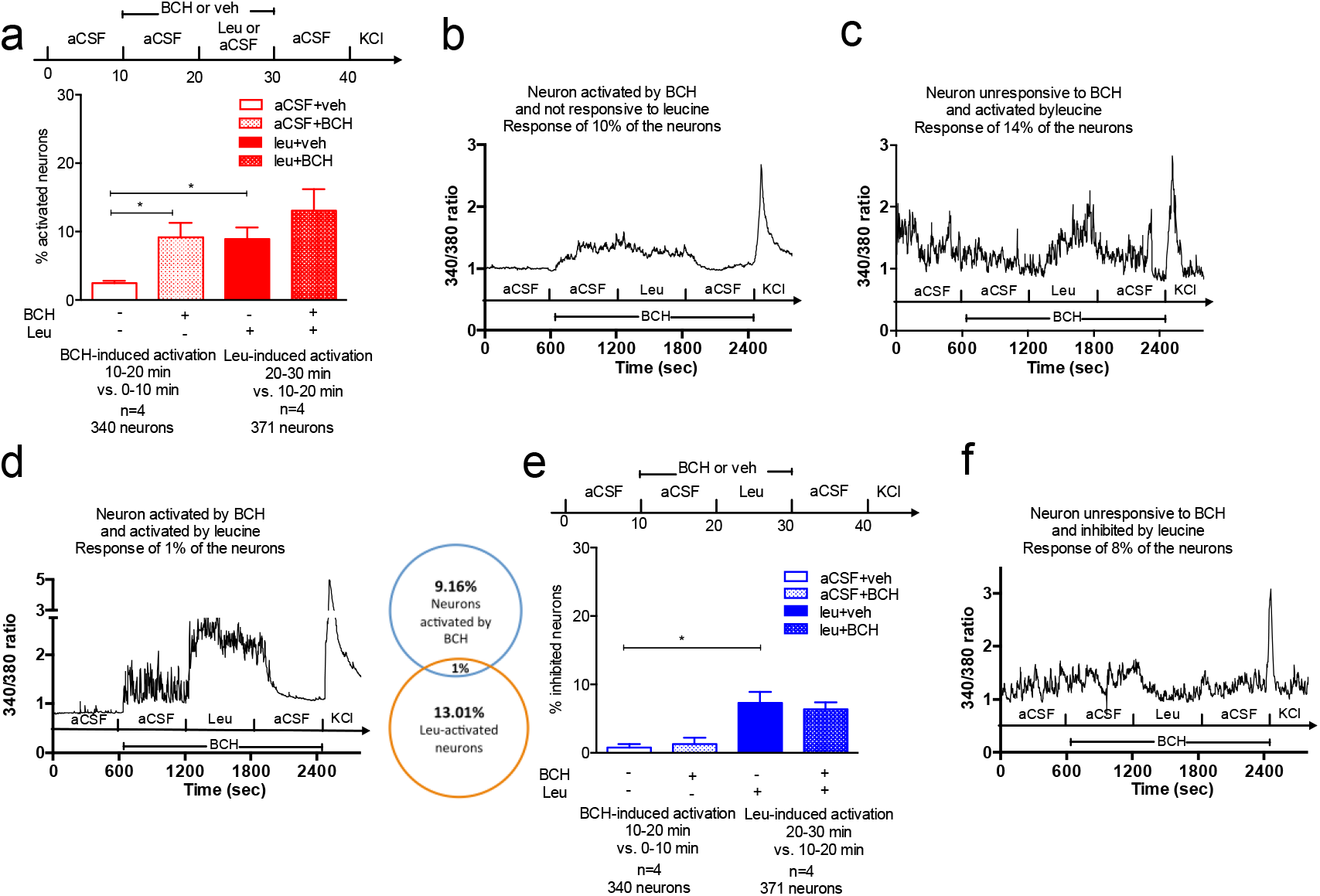
Leucine-induced changes in [Ca2^+^]_i_ rely on extracellular leucine detection. Neuronal responses to leucine in the presence of (a, e) in primary cultures of mediobasal hypothalamic neurons. Changes in [Ca^2+^]_i_ in primary cultures of mediobasal hypothalamic neurons treated with leucine and BCH (b, c, d, f). Data are means ± SEM. *:p<0.05.

### 3.7 Mechanistic insights into leucine-induced calcium fluxes

In neurons, calcium released from endoplasmic reticulum (ER) stores or calcium entering the cell through voltage or ligand gated ion channels are the principal sources that contribute to changes in intracellular calcium concentrations in response to a ligand. We used multiple strategies to determine which source of calcium is mobilized by leucine to generate changes in [Ca^2+^]_i_.

We first tested the role of intracellular calcium stores using thapsigargin, an irreversible non-competitive inhibitor of the SERCA ATP_ase_, that pumps calcium into the endoplasmic reticulum against its concentration gradient, either to compensate for inward calcium leakage, or to refill the stores after emptying in response to intracellular signaling events (38). Cells were pretreated with 200 nM thapsigargin for 10 minutes to deplete intracellular calcium stores (38) and then treated with leucine or vehicle in the presence of Thapsigargin. Upon exposure to Thapsigargin, 66.4% cells showed a calcium response confirming the depletion of intracellular calcium stores (Figure 6a). After this initial rise, baseline [Ca^2+^]_i_ remained elevated in 25% of cells, with a Fura2 340/380 ratio on average 0.4 higher than during aCSF exposure (Figure 6a). In the presence of thapsigargin, leucine was still able to produce an increase in [Ca^2+^]_i_ in 17% of cells (Figure 6b, 6c). By contrast, leucine produced inhibition in a marginal number of cells in the presence of Thapsigargin (Figure 6c).

**Figure 6:**
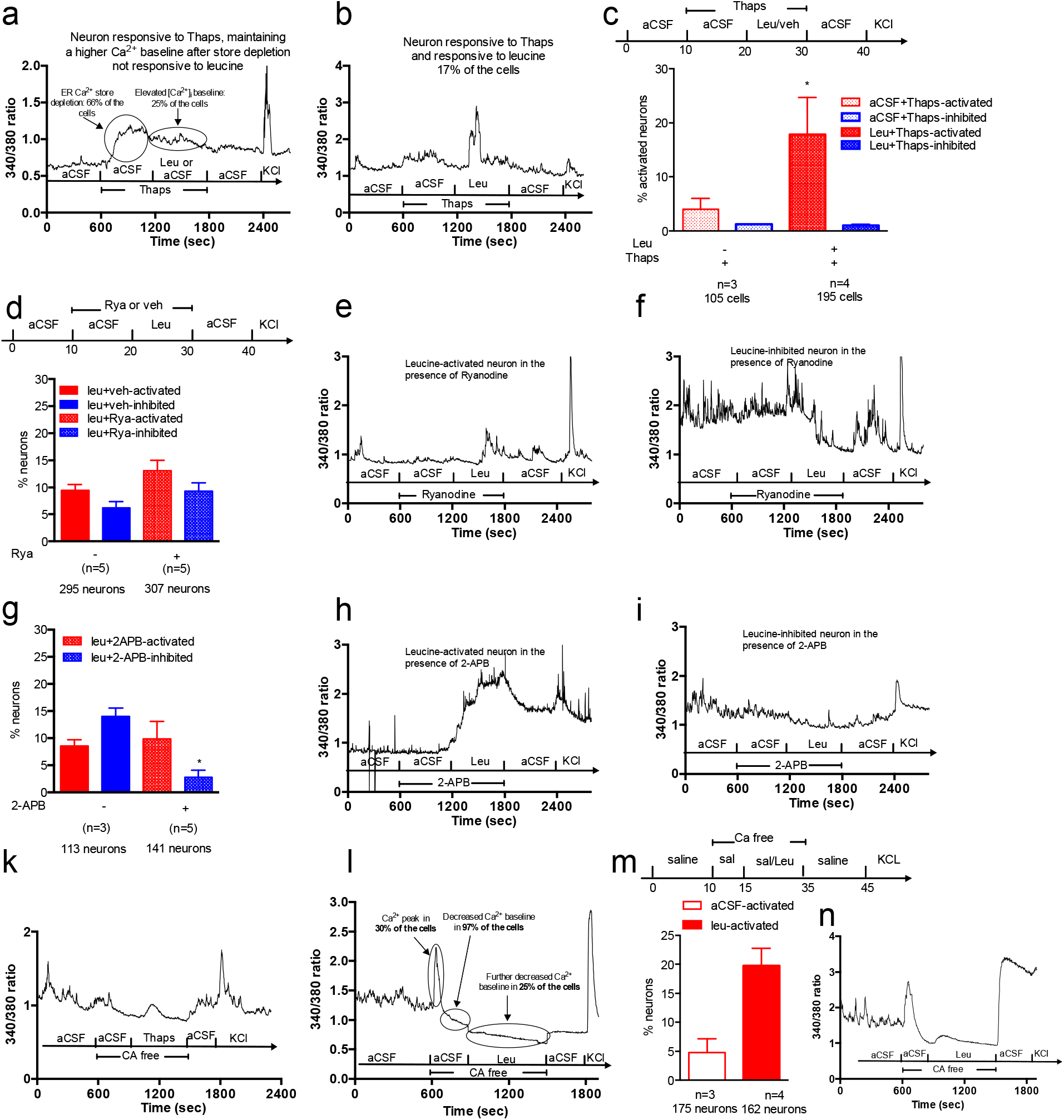
Role of calcium channels in hypothalamic leucine sensing. Traces of a typical response of neurons in culture to thapsigargin exposure (a). Trace of a leucine-activated neuron in the presence of thapsigargin (b). Percentage of leucine-activated and leucine-inhibited neurons in the presence of thapsigargin (c). Percentage of leucine-activated and inhibited neurons in the presence of Ryanodine (d) and traces of a leucine-activated (e) and a leucine-inhibited (f) neuron in the presence of ryanodine. Percentage of leucine-activated and inhibited neurons in the presence of 2-APB (g). Trace of a leucine-activated neuron in the presence of 2-APB (h). Trace of a leucine-inhibited neurons in the presence of 2-APB (i). showing the effect of thapsigargin on calcium concentration following a 5 minutes treatment in calcium depleted media (j). Trace showing the response to the calcium depleted media in neurons with a calcium leak (k). Percentage of leucine-activated neurons in calcium depleted conditions (l) and trace of a leucine-activated neurons in calcium depleted conditions (m). Data are means ± SEM. *:p<0.05.

We then examined the specific channels involved in the release of calcium from the endoplasmic reticulum, and blocked ryanodine receptors (RyR) with ryanodine, which binds and rapidly induces closure of RyR at concentrations above 100nM (39, 40). Co-application of 25 µM ryanodine with leucine did not significantly change the percentage of cells activated or inhibited by leucine (Figure 6d-6f). In POMC neurons, ryanodine also did not change the percentage of cells activated by leucine (20% cells activated with DMSO+Leu, n=5, 21 POMC neurons; 20% cells activated with Rya+Leu, n=5, 26 POMC neurons). Likewise, 100 µM 2-APB, an inhibitor of IP_3_R (41) did not block leucine-induced activation (Figure 6g, 6h), but significantly lowered the percentage of leucine-inhibited cells (Figure 6g, 6i). Thus, leucine-induced neuronal activation does not rely on RyR and IP_3_R channels, while leucine-induced inhibition is mediated at least in part by a 2-APB sensitive mechanism.

Last, to test whether leucine-induced activation requires calcium entry from across the plasma membrane, responses to leucine were measured in cells pre-incubated with a calcium-depleted aCSF media for 5 minutes. Under these conditions, transmembrane calcium fluxes are abrogated, but ER stores remain intact, as confirmed by the ability of 200nM thapsigargin to produce a rapid transient increase in intracellular calcium concentrations (Figure 6k). Upon transition to the calcium depleted media, 30% of cells showed a brief calcium peak (Figure 6l), and 96.8% of neurons showed a rapid decrease in intracellular calcium levels, reaching a stable lowered baseline within 5 minutes. However, in 25% of cells, [Ca^2+^]_i_ further decreased during the following 10 minutes, reaching a nadir Fura 2 340/380 ratio of 0.5–0.6 (Figure 6l). Following a 5 minutes treatment with a calcium depleted media, the percentage of neurons activated by leucine remained the same (Figure 6m) but the magnitude of the calcium response was significantly blunted (increase in Fura2 340/380 over baseline of 0.09±0.029 vs. 0.63±0.031, p<0.01), and the remaining Ca^2+^ fluxes were more transient than those measured in the presence of extracellular calcium (Figure 6n). In POMC neurons, such a small response to leucine during incubation with a calcium depleted media was observed in only 4% of neurons (n=7, 23 POMC neurons). By contrast, incubation in a calcium-depleted media abrogated all leucine-induced neuronal inhibition.

Collectively, these data indicate that leucine-induced activation of hypothalamic neurons principally relies on calcium entry from the extracellular calcium pool. The mechanisms underlying leucine-induced inhibition of hypothalamic neurons are sensitive to both SERCA ATP_ase_ inhibition and 2-APB.

### 3.7 hPSC-derived hypothalamic neurons sense leucine

The studies performed in acutely dissociated mouse MBH neurons suggested that leucine is sensed by a subset of MBH neurons that respond by altering their activity and neuropeptide secretion to regulate energy homeostasis. Since mice and humans show similar behavioral responses to dietary amino acids (8), we hypothesized that human hypothalamic neurons also show rapid responses to leucine. To test this hypothesis, we differentiated human pluripotent stem cells into hypothalamic neurons as previously described [Merkle, 2015 #636] [Kirwan, 2017 #471]. After at least 30 days in culture, these *in vitro*-derived human hypothalamic cultures contained many neurons that were strongly immunopositive for POMC, suggesting they had been patterned to a BMH-like regional identity [Kirwan, 2017 #471]. After cells had matured at least 30 days in culture, cultures were dissociated, re-plated, and imaged 24 hours later after loading with Fura2 dye to replicate the experimental manipulations that primary murine MBH neurons had undergone. We found that a subset of human hypothalamic neuron cultures responded to 500 uM leucine. Using identical criteria to define leucine-activated and leucine-inhibited cells as in experiments performed with murine culture, we found that 10.4±3.2 % of hPSCs-derived hypothalamic neurons (n=6, 212 neurons) increased their [Ca^2+^]_i_ in response to leucine (Figure 7a), producing a peak in [Ca^2+^]_i_ that was 25% greater than in vehicle treated cells and an increase in the Fura2 340/380 AUC of 60% (Figure 7b-7c). Leucine decreased [Ca^2+^]_i_ in 15.6±7.2 % of hPSCs-derived hypothalamic neurons (n=6, 212 neurons) and produced an average 9% decrease in the minimum Fura2 340/380 ratio and a 55% decrease in the Fura2 340/380 AUC (Figure 7d-7f). These results are similar to those observed in murine MBH neurons (Fig. 2).

**Figure 7:**
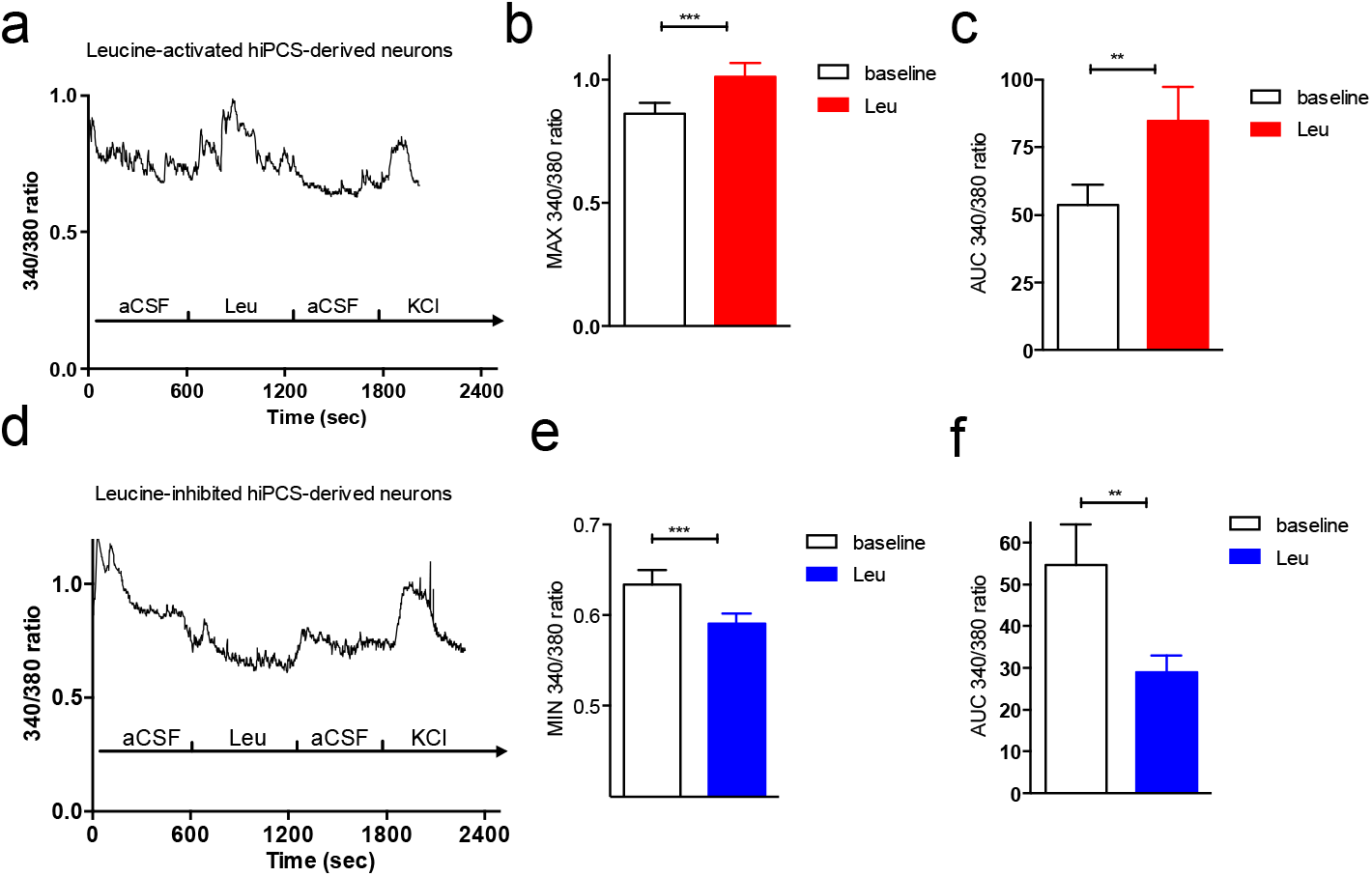
Leucine sensing in hypothalamic neurons derived from human pluripotent stem cells. 340/380 nm fluorescence (a), peak 340/380 ratio (b) and 340/380 ratio AUC (c) in human hypothalamic neurons activated in response to 500 µM leucine exposure. 340/380 nm fluorescence (d), minimum 340/380 ratio (e) and 340/380 ratio AUC (f) in human hypothalamic neurons inhibited in response to 500 µM leucine exposure. Data are mean ± SEM. *: p<0.05. **: p<0.01. ***: p<0.001

## 4. Discussion

We developed a model of primary culture of dissociated mouse MBH neurons prepared from post-weaning mice to investigate cell autonomous nutrient-sensing properties of mature MBH neurons. Using this model, we found that leucine sensing is not a ubiquitous property of MBH neurons. Instead, only a subset of MBH neurons responded to a physiologically relevant increase in the extracellular concentration of leucine. Leucine produced heterogeneous responses in a neurochemically diverse group of MBH neurons including a subset of POMC and NPY/AGRP neurons. Using a series of pharmacological manipulations, we gained insights into the mechanisms engaged by leucine to rapidly alter the activity of MBH neurons. Unexpectedly, pharmacological gain- or loss-of-function of pathways previously implicated in hypothalamic leucine sensing failed to alter the rapid calcium responses to leucine. Instead, our data implicate activation of plasma membrane Ca^2+^ channels in leucine-induced neuronal activation, and a mechanism sensitive to SERCA ATP_ase_ inhibition and 2-APB in leucine-induced neuronal inhibition. Last, we found that a subset of iPSC-derived human hypothalamic neurons responded to leucine and these responses showed remarkable similarity to those obtained in murine cultures, suggesting that the data obtained with primary mouse neurons have direct relevance to human energy homeostasis.

Our data confirm that ARH POMC neurons represent one of the main MBH neuronal population activated by leucine (10, 42). Although melanocortinergic signaling has been implicated in the feeding response to MBH Leu, POMC neurons are unlikely to mediate the rapid behavioral response to Leu, as optogenetic or chemogenetic activation of ARH POMC neurons reduce food intake only hours after stimulation (43, 44). How rapid changes in POMC neuronal excitability affect appetite over that time frame remains to be determined. Alternatively, this rapid leucine sensing may regulate other physiological functions downstream of POMC-regulated neural circuits, including glucose homeostasis. By contrast, rapid inhibition of a subset of NPY/AGRP neurons is likely to mediate the rapid increase in first meal latency and suppression of energy intake measured following acute leucine nano-injection into the MBH (10), as acute inhibition of AGRP neurons rapidly suppresses appetite in hungry mice (45–47). Consistent with such an effect, we show that leucine can suppress AGRP secretion from hypothalamic slices of fasted mice. These results represent the first piece of evidence showing that amino acids can rapidly inhibit one of the most critical orexigenic neuronal population of the hypothalamus, and provide a mechanism for the rapid inhibition of foraging behaviour in rodents following parenchymal administration of leucine into the MBH (10). Previous work showed that leucine can regulate AGRP mRNA expression in GT1–7 cells, indicating that Leucine sensing may also produce long term consequences on AGRP neuronal output (48). Taken together, our result showing both activation of ARH POMC neurons and inhibition of ARH AGRP/NPY neurons are consistent with the anorectic consequences of central leucine sensing, and reveal that the multimodal anorectic behavioural response to MBH leucine exposure (i.e. increased first meal latency, decreased meal size, decreased meal frequency over 24h) is mediated via the modulation of the activity of multiple neuronal populations and downstream neural circuits.

One of the key findings of our study is that pathways previously implicated in hypothalamic leucine sensing – mTORC1 signaling, leucine decarboxylation, and K_ATP_ channels – are not required for the rapid changes in intracellular Ca^2+^ concentrations of MBH neurons in response to leucine. Consistent with these results, genetic models developed so far to target these pathways failed to produce convincing evidence implicating them in long-term body weight control (42, 49). Additional evidence clearly indicate that leucine can signal independently of mTORC1 to alter the activity of hypothalamic neurons. Lack of the mTORC1 effector rpS6k1 in POMC neurons failed abrogate the ability of ARH POMC neurons to depolarize in response to leucine (42), and rapamycin failed suppress the effect of hypothalamic leucine on hepatic glucose output (50). Our data suggest that activation of mTORC1 signaling may not be essential for the rapid appetitive and consumatory inhibitory behavioural responses to hypothalamic leucine. This does not preclude a role for mTORC1 signaling in the control of food intake, as rapamycin can rapidly produce hunger in sated animals (50, 51).

Leucine has been proposed to signal in the hypothalamus in part via products of its metabolism feeding into the TCA cycle and leading to ATP production and modulation of the activity of K_ATP_ channels (10, 50). Our results indicate that rapid leucine sensing by MBH neurons does not require leucine catabolism, as leucine’s ketoacid did not modulate Ca^2+^ concentrations of MBH neurons in culture. This observation is consistent with the lack of an acute anorectic response to parenchymal KIC administration in rodents (10). In addition, we show that functional K_ATP_ channels are not required for the rapid response to leucine. In fact, our data indicate that most leucine sensing neurons do not respond to tolbutamine, diazoxide or BCH, and thus, do not appear to have functional K_ATP_ currents. Although diazoxide has been shown to rapidly open K_ATP_ channels (52, 53) the paradigm we used in this particular experiment with simultaneous exposure to diazoxide and leucine does not allow us to make strong conclusions. However, together with the results of the other approaches we used, we can conclude that leucine-induced activation does not require functional K_ATP_ channels, and that metabolic sensing neurons relying on the modulation of the activity K_ATP_ channels do not markedly overlap with the neuronal populations that rapidly sense changes in leucine concentrations. This conclusion raises the question of the potential segregation between leucine and glucose sensing neurons, and supports the idea that leucine-sensing neurons represent a neuronal population distinct from the glucose-sensing neurons that have functional and glucose-sensitive K_ATP_ channels.

Instead, our findings suggest that neuronal leucine sensing requires Ca^2+^ influx through the plasma membrane, as responses were severely abrogated in Ca^2+^ depleted media. Leucine-induced activation appeared to be insensitive to manipulations of ER calcium stores, as shown by the lack of effect of thapsigargin, ryanodine and 2-APB pretreatment on leucine-induced calcium elevations. The insensitivity to 2-APB also excludes a possible contribution of TRPC channels that open upon calcium release from intracellular stores [Bavencoffe, 2017 #1257], but are 2-APB sensitive. These results also support the conclusion that leucine is sensed extracellularly, at least in leucine-activated neurons. Consistent with this conclusion, we showed that rapid neuronal leucine sensing is independent of system L transporters. However, other transporters mediate the uptake of leucine into neurons, in particular slc6a15, slc6a16 and slc6 a17 (54, 55), expressed in mouse neurons [Zhang, 2014 #1268], BCH can potently inhibit slc6a15 (85% inhibition with 10mM BCH) (55, 56), but the effects of BCH on slc6a16 and slc6a17 have not been characterized.

Our data do not allow us to draw strong conclusions about the mechanisms underlying leucine-induced inhibition. The sensitivity of leucine-induced inhibition to both thapsigargin and 2-APB raises the possibility that leucine-inhibited neurons have a constitutive leaking inward calcium current responsive to depletion of ER calcium stores. This is supported by the ability of thapsigargin to increase baseline [Ca^2+^]_i_ in 30% of cells, and the inability of 24% of cells to maintain a stable baseline [Ca^2+^]_i_ when exposed to a calcium-depleted media. This calcium leak could come from a store-operated calcium channel, and a conclusion consistent with all our observation would be that in leucine-inhibited neurons, leucine either directly or indirectly inhibits a store operated calcium current, leading to decreased [Ca^2+^]_i_. This hypothesis will have to be directly tested in the future. Of note, the mechanisms underlying glucose-induce d neuronal inhibition are also not clarified and are still under debate.

Recently, hypothalamic tanycytes were shown to sense amino acids from the ventricular compartment via Tas1r1/Tas1r3 and mGluR4 [Lazutkaite, 2017 #1258], Here, we can exclude a contribution of these receptors that function through G-protein mediated opening of IP_3_R, which we show are not required for leucine-induced activation of hypothalamic neurons.

## 5. Conclusion

This work represents an important conceptual advance in the characterization and understanding of how hypothalamic circuits sense amino acid and protein availability. We showed that a specialized group of MBH neurons rapidly respond to physiological changes in extracellular leucine concentrations. The leucine-sensing neuronal population of the MBH overlaps with known neurochemical populations of this area. In addition, leucine can produce a variety of responses, supporting the conclusions that leucine can be sensed by distinct neural circuits via different mechanisms, to control a variety of downstream functions. Neuronal leucine sensing in the mediobasal hypothalamus does not rely on mTORC1 signaling, leucine metabolism or K_ATP_ channels, but requires the modulation of plasma membrane calcium channels (Summary figure). Future work is required to better characterize these mechanisms and identify the unique molecular components that make a hypothalamic neuron responsive to leucine. Such work is critical for the future in vivo manipulation of leucine-sensing pathways and to test their relevance to body weight control and metabolic function.

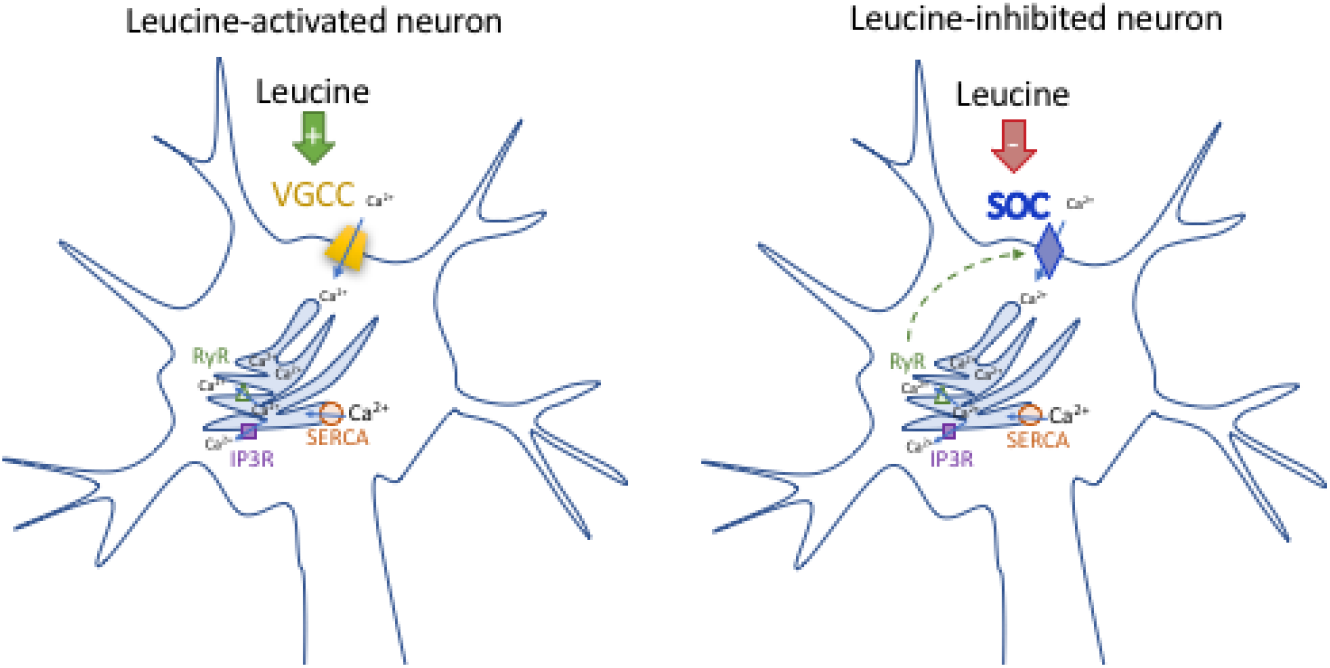
Summary Figure: Proposed mechanisms implicated in rapid leucine sensing in mediobasal hypothalamic neurons. Leucine rapidly modulates the activity of a subset of mediobasal hypothalamic neurons.. In leucine-activated neurons (including POMC neurons) leucine activates a voltage-gated calcium channel (VGCC). In leucine-inhibited neurons, leucine inhibits a store-operated calcium current (SOC).

## Acknowledgments

This work was supported by the Medical Research Council New Blood Fellowship [MR/M501736/1] to CB, the Medical Research Council Metabolic Disease Unit [MRC_MC_UU_12012/5], and the Wellcome Trust Strategic award for the MRL Disease Model Core and Imaging facilities [100574/Z/12/Z].

## Footnotes

1 Abbreviations: ARH: arcuate nucleus of the hypothalamus, MBH: mediobasal-hypothalamus, hPSC: human induced pluripotent stem cells,

